# The SARS-CoV-2 exerts a distinctive strategy for interacting with the ACE2 human receptor

**DOI:** 10.1101/2020.03.10.986398

**Authors:** Esther S. Brielle, Dina Schneidman-Duhovny, Michal Linial

**Affiliations:** Department of Biological Chemistry, Institute of Life Sciences, The Hebrew University of Jerusalem, Jerusalem, Israel; The Alexander Grass Center for Bioengineering, The Hebrew University of Jerusalem, Jerusalem, Israel; The Rachel and Selim Benin School of Computer Science and Engineering, The Hebrew University of Jerusalem, Jerusalem, Israel

## Abstract

The COVID-19 disease has plagued over 110 countries and has resulted in over 4,000 deaths within 10 weeks. We compare the interaction between the human ACE2 receptor and the SARS-CoV-2 spike protein with that of other pathogenic coronaviruses using molecular dynamics simulations. SARS-CoV, SARS-CoV-2, and HCoV-NL63 recognize ACE2 as the natural receptor but present a distinct binding interface to ACE2 and a different network of residue-residue contacts. SARS-CoV and SARS-CoV-2 have comparable binding affinities achieved by balancing energetics and dynamics. The SARS-CoV-2–ACE2 complex contains a higher number of contacts, a larger interface area, and decreased interface residue fluctuations relative to SARS-CoV. These findings expose an exceptional evolutionary exploration exerted by coronaviruses toward host recognition. We postulate that the versatility of cell receptor binding strategies has immediate implications on therapeutic strategies.

**One Sentence Summary:** Molecular dynamics simulations reveal a temporal dimension of coronaviruses interactions with the host receptor.

## Main Text

Coronavirus disease 2019 (COVID-19), initially detected in the Wuhan seafood market in the Hubei province of China (*1*) is caused by SARS-CoV-2 (referred to as the COVID-19 virus for clarity). The COVID-19 virus already spread within 10 weeks from its appearance to more than 110 countries, resulting in over 4,000 deaths worldwide. The COVID-19 virus is capable of human-to-human transmission and was introduced to humans in a zoonotic event (*2*).

Currently, seven confirmed coronavirus (CoV) species are known as human pathogens (*3*). Four CoV are endemic species in humans and cause mild respiratory symptoms, mostly in pediatric patients (*4*). These are the HCoV-HKU1 and HCoV-OC43 from the betacoronavirus (BCoV) genus and the HCoV-229E, and HCoV-NL63 from the alphacoronavirus (ACoV) genus. The other human CoVs have caused severe outbreaks (**Table S1**). The SARS-CoV (referred to as the SARS-2002 virus for clarity), is a BCoV that emerged in humans in 2002, giving rise to the Severe Acute Respiratory Syndrome (SARS) outbreak. The Middle East Respiratory Syndrome (MERS) BCoV caused an outbreak in 2012-2013. Most recently, SARS-CoV-2, with high homology to the 2002 SARS-CoV, caused the current pandemic-like COVID-19 outbreak (*5*).

To gain access to host cells, coronaviruses rely on spike proteins, which are membrane-anchored trimers containing a receptor-binding S1 segment and a membrane-fusion S2 segment (*6*). The S1 segment contains a receptor binding domain (RBD) that recognizes and binds to a host cell receptor. The angiotensin-converting enzyme 2 (ACE2) was identified as the critical receptor for mediating SARS-2002 entry into host cells (*7, 8*). Binding of the spike protein to the receptor is a critical phase where the levels of the ACE2 expressed on the cell membrane correlates with viral infectivity, and govern clinical outcomes (*9*). Consistent with the clinical pulmonary manifestation, ACE2 is widely expressed in almost all tissues, with the highest expression levels in the epithelium of the lung (*10*). Similar to the SARS-2002 virus, the COVID-19 virus enters the host cell by RBD binding to the host cell ACE2 receptor (*7, 11, 12*). Host receptor recognition for cell entry is, however, not specified by the CoV genus classification. MERS-CoV is a member of the BCoV genus but does not recognize the ACE2 receptor. In contrast, HCoV-NL63 is a member of the ACoV genus and does recognize the ACE2 receptor (*13*).

Herein, we analyze the binding of several CoV RBDs to ACE2 with molecular dynamics (MD) simulations and compare the stability, relative interaction strength, and dynamics of the interaction between the viral spike protein and the human ACE2 receptor.

The COVID-19 RBD (residues 319-529) shares a 72.8% sequence identity and high structural similarity with the SARS-2002 RBD (**Table 1**). In contrast, the RBD of HCoV-NL63 is only 17.1% identical to that of COVID-19 and there are no significant structural similarities between them (**Fig. S1**). Remarkably, the RBD of MERS-CoV, which is structurally similar to that of COVID-19 (20.1% sequence identity, 65% structure similarity) recognizes a different host receptor (DPP4) for its cell entry and does not bind ACE2 (*14*).

**Table 1:**
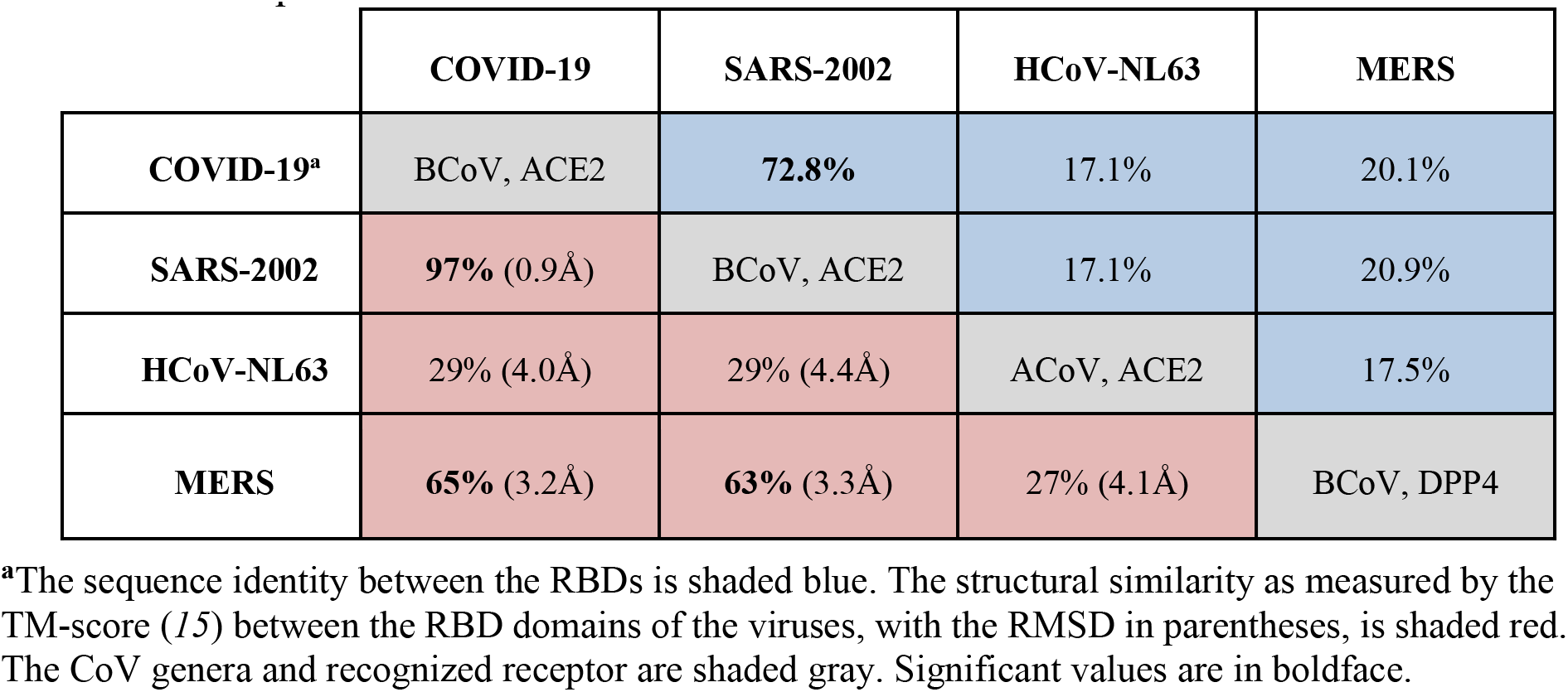
The sequence and structural resemblance of the RBDs of various human coronaviruses.

We ran 100ns molecular dynamic (MD) simulations of ACE2 in complex with the RBDs of the COVID-19, SARS-2002, and HCoV-NL63 viruses to quantify the energetics and the dynamics of the different RBD—ACE2 interactions. The simulation trajectory snapshots at 10 ps intervals (10,000 frames) were analyzed by a statistical potential to assess the probability of the RBD— ACE2 interaction (SOAP score, (*16*)), with lower values corresponding to higher probabilities and thus higher affinities. The interaction scores for COVID-19 RBD—ACE2 were comparable to those of SARS-2002, median of −1865.9 and −1929.5, respectively (**Fig. 1A**). HCoV-NL63 has RBD—ACE2 interaction scores are higher than both of the SARS-CoVs (median of −941.6). MERS, which is structurally similar to COVID-19 (**Table 1**) does not bind ACE2. MERS virus which binds dipeptidyl peptidase-4 (DPP4, also known as CD26 (*14*)), has RBD—ACE2 interaction scores that indicate extremely weak affinity (median of −692.6), as expected from a non-cognate receptor interaction. COVID-19 has the largest buried surface area at the interface (1204Å^2^), followed by the interface area for SARS-2002 (998Å^2^) and HCoV-NL63 (973Å^2^). The number of ACE2 contacting residues maintains the same order, with 30, 24, and 23 for COVID-19, SARS-2002, and HCoV-NL63, respectively (**Fig. 1C**). The three RBDs exploit specific binding sites on ACE2 based on the analysis of the MD trajectories (**Fig. 1, C and D**; **Movie S1**). There is a significant overlap of ACE2 interacting residues between COVID-19 and SARS-2002 (at least 73%), while HCoV-NL63 shares only 17% and 36% of contacts with SARS-2002 and COVID-19, respectively. These findings suggest that the coronaviruses exert different interaction strategies with their cognate receptors to achieve the affinity that is required for effective cell entry.

**Fig. 1.**
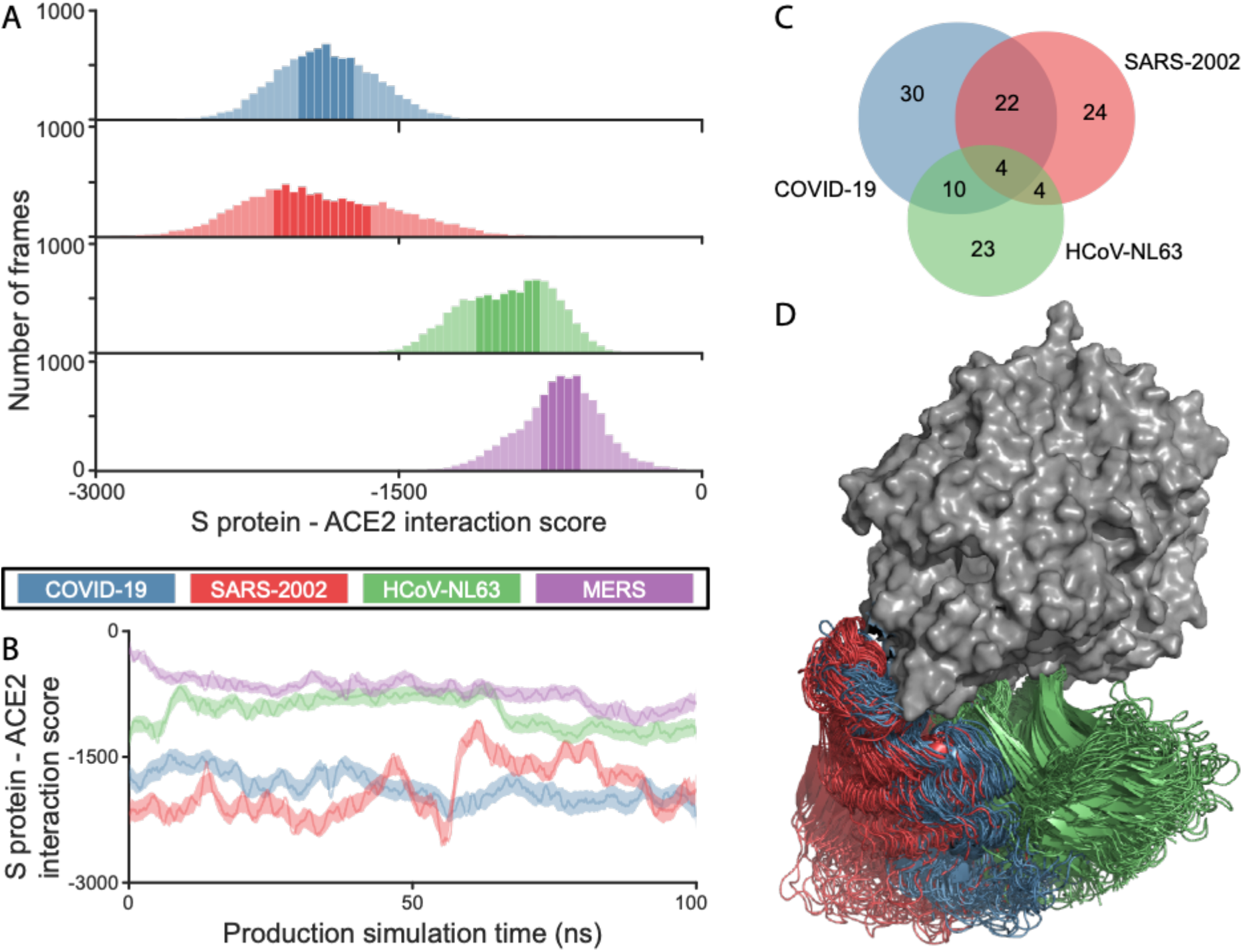
Analysis of RBD—ACE2 interactions based on MD trajectories. COVID-19, SARS-2002, and HCoV-NL63 are colored blue, red, and green, respectively. **(A)** Histograms of the RBD—ACE2 interaction scores throughout the simulation trajectory. Darker color represents 75% of all frames. **(B)** The score values along the simulation trajectory, smoothed along the elapsed time. **(C)** Venn diagram of ACE2 interacting residues for COVID-19, SARS-2002, and HCoV-NL63. An ACE2 residue is considered as part of the interface if one of its atoms is within 4Å from any RBD atom in at least 10% of the 10,000 MD simulation frames. **(D)** Overlay of 50 snapshots for each of the three RBDs. The ACE2 is in surface representation (gray). The frames were aligned using the N-terminal fragment of ACE2 that contains the two helices participating in the RBDs binding.

While the sequence identity between the RBDs of COVID-19 and SARS-2002 is 73% (**Table 1**), we observe a significantly higher residue substitution rate at the interaction interface with the ACE2 receptor. Out of 29 RBD interface residues, only 10 residues (34%) in COVID-19 are conserved with respect to SARS-2002 (**Fig. 2A, Table S1**, **Fig. S1**). Similarly, only 12 residues (40%) in SARS-2002 are conserved with respect to COVID-19.

**Fig. 2:**
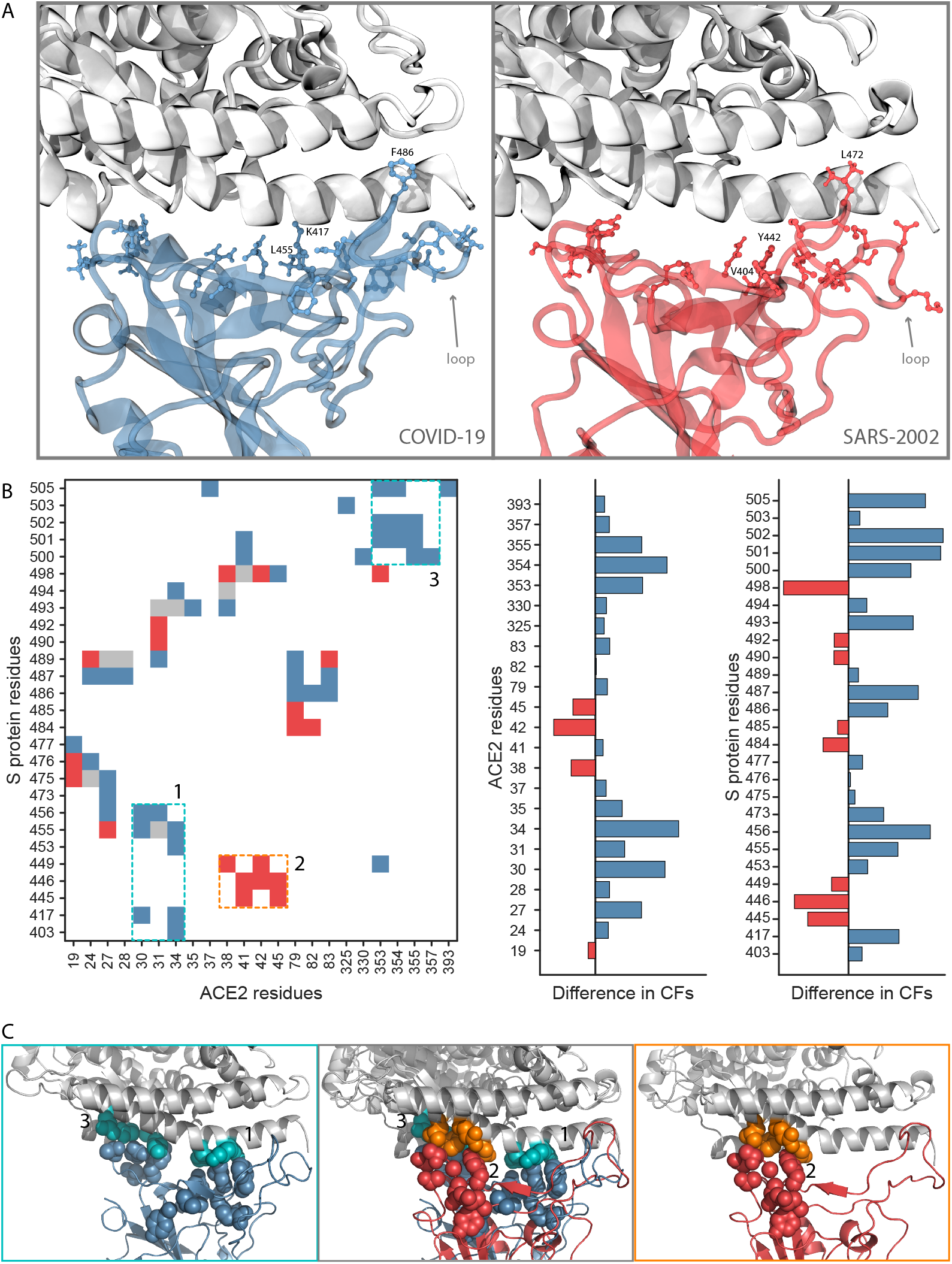
Interaction interfaces of RBD—ACE2. A residue is considered as part of the interface if one of its atoms is within 4Å from any atom of the other partner in at least 30% of the 10,000 MD simulation frames. **(A)** Interface residue side-chain heavy atoms that vary between COVID-19 (blue) and SARS-2002 (red) are shown with ball-and-stick representations. ACE2 is colored gray. **(B)** Contacts difference plot between COVID-19 and SARS-2002. Contacts with 50% greater CF in COVID-19 RBD—ACE2 vs. SARS-2002 RBD—ACE2 are colored blue. Contacts with 50% greater CF in SARS-2002 RBD—ACE2 vs. COVID-19 RBD—ACE2 are colored red. Similar interface-residue CFs (<50%) in both RBDs are colored gray. The residue numbering is according to COVID-19 (RefSeq: YP_009724390.1). **(C)** Difference plots for interface residue CFs that are in the interface for ACE2 (left) and RBD (right). **(D)** Zoom on the interface contacts unique to each virus.

To investigate these interface residues, we construct and overlay the contact maps for the RBD— ACE2 interfaces for COVID-19 and SARS-2002 (**Fig. 2B**). We define a residue-residue contact frequency (CF) as the fraction of MD trajectory frames in which the contact appears. Remarkably, only 8 out of the total 72 residue-residue interface contacts have comparable (<50% difference) contact frequencies between the COVID-19—ACE2 and SARS-2002—ACE2 interfaces (**Fig. 2B**, colored gray). Furthermore, we find two interaction patches unique to COVID-19 (**Fig. 2B**, patches 1 and 3) and another patch unique to SARS-2002 (**Fig. 2B**, patch 2). COVID-19 has a significant and unique contact site between residues 500-505 of the RBD and residues 353-357 of ACE2 (**Fig. 2, B and C**). COVID-19 also creates a new interaction patch with the middle of the N-terminal ACE2 helix (**Fig. 2, B and C)**, while SARS-2002 has a unique interaction patch with the end of the same helix (**Fig. 2, B and C)**. The rest of the changes in the interface contact frequencies are due to the different interface loop conformations (COVID-19 residue numbers 474-498, SARS-2002 residue numbers 461-484) (**Fig. 2, A and B, Table S1**). COVID-19 has a significantly higher number of well-defined contact pairs compared to SARS-2002: 52 vs. 28 contacts (with 44 and 20 unique pairs, excluding the ones with similar CFs) were found for RBD—ACE2 of the COVID-19 and SARS-2002, respectively (**Fig. 2B**). Results from Fig. 2 expose the accelerated evolution among the key anchoring residues of the RBD—ACE2 interface. This comparison raises the following question: How does SARS-2002 RBD reach an ACE2 binding affinity that is comparable to that of COVID-19 but with fewer contact pairs and a smaller interface area?

The distribution of SOAP scores throughout the simulation trajectory has a larger fluctuation range for SARS-2002, relative to COVID-19 (**Fig. 1, A and B**; **Fig. S2A)** suggesting that SARS-2002—ACE2 interaction is fluctuating between several structural states. Moreover, analysis of contact frequencies along the entire trajectory reveals that none of the SARS-2002 contacts are maintained over 90% of the frames while COVID-19 still maintains about half of its contacts at 90% of the trajectory (**Fig. S2B**).

To investigate the dynamics of COVID-19 binding compared to SARS-2002, we calculate the root-mean-square fluctuation (RMSF) of each residue with respect to the lowest energy snapshot from their respective 100ns MD simulation trajectory. The interface region in the RBD contains two loops (loop1: residues 474-489, loop2: residues 498-505; using COVID-19 numbering, **Fig. 3D**) that bind to the ACE2 N-terminal helix on both of its ends. These two loops are highly flexible in the SARS-2002 RBD (**Fig. 3, A and D**). While loop1 is also fluctuating in the COVID-19 RBD, albeit much less, loop2 remains relatively rigid in the COVID-19 RBD. In addition, we find that in the COVID-19-RBD, a region centered around K417 leads to further stability relative to the corresponding region in SARS-2002. We attribute this difference to the unique interaction of COVID-19 at position K417 with the middle of the N-terminal ACE2 helix, thus serving as an anchor site to the receptor (**Fig. 2C** and **Fig. 3A**). The contribution of K417 to ACE2 binding is observed in a recent cryoEM structure of the COVID-19 spike protein bound to ACE2 (*17*). Overall, COVID-19 is more rigid compared to SARS-2002 (**Fig. 3, A and D**).

**Fig. 3.**
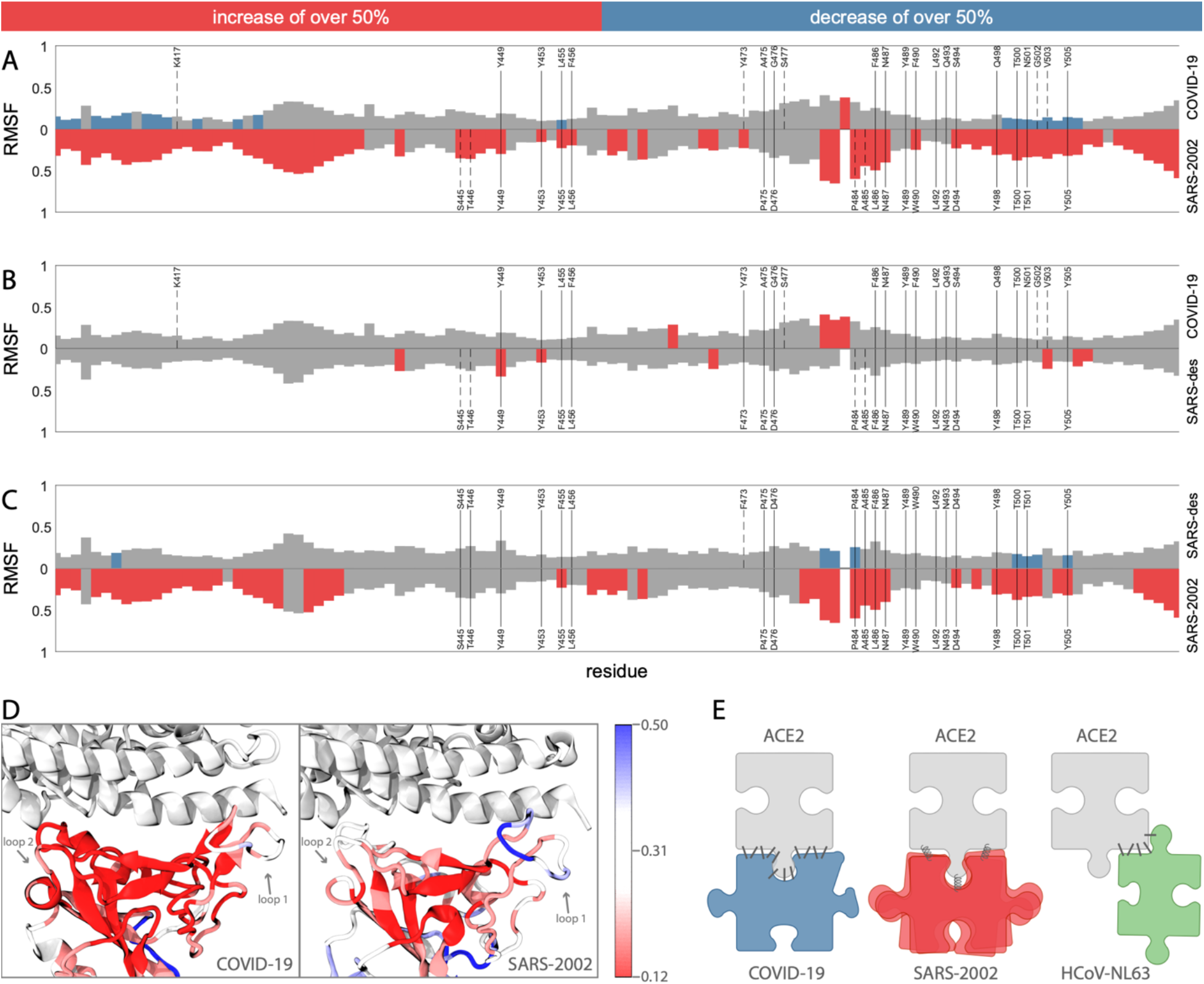
Dynamics of the RBD domains. The dynamics of COVID-19 and SARS2002, with a comparison to a designed SARS mutant are shown. The graphs show the root-mean-square fluctuation (RMSF) of each residue along the simulation trajectory with respect to the structure with minimum energy. The residue numbers are according to COVID-19 numbering (RefSeq: YP_009724390.1). The top graph **(A)** compares COVID-19 with SARS-2002. The middle graph **(B)** compares COVID-19 with SARS-designed. The bottom graph **(C)** compares SARS-2002 with SARS-designed. For all three graphs, positions highlighted in red or blue indicate those that have an increase or decrease, respectively, of 50% RMSF with respect to the comparison graph. Contact positions are written in gray, with solid vertical lines denoting the contact residues that exist in both comparison structures, and dashed vertical lines denote contact residues that exist in only one of the comparison structures.

We investigate the dynamics of a designed SARS (SARS-des) variant (*11*), which differs from SARS-2002 at only 2 positions: Y455F and L486F (The matched position in SARS-2002 residues are 442 and 472, respectively; **Table S1**). The L486F mutation is of special interest for the COVID-19 RBD as well because it has this same substitution. Our MD simulation analysis reveals that the SARS-des has a substantially lower interaction scores with ACE2 (median of −2199.2, **Fig. S2**), as expected for an optimized human ACE2-binding RBD design. We observed that these two mutations not only enhance the binding affinity to ACE2, but also lead to a substantial stabilization of the interaction interface. The fluctuation signatures along the RBD of SARS-des are surprisingly similar to those recorded for COVID-19 (**Fig. 3, B and C**). Thus, the switch from a flexible binding mode (for SARS-2002) to a stable one (COVID-19 and SARS-des, **Fig. 3B**) highlights the remarkable capacity of the RBD to adopt alternative receptor binding strategies driven by a minimal number of amino acid substitutions. This analysis reveals the critical role of L486F (SARS-des residue F472) for stabilizing the COVID-19—ACE2 interface and a reduction in the number of states of the COVID-19 spike protein bound to an ACE2 receptor.

Experimental affinity measurements (e.g. surface plasmon resonance, SPR) confirm the high affinity of SARS-2002 RBD—ACE2 binding, with an equilibrium dissociation constant (K_D_) of ~15 mM (*18–21*), similar to the binding affinity of ACE2 and the COVID-19 RBD (*22, 23*). Our MD based calculation is consistent with SARS-2002 displaying a similar but slightly higher affinity relative to COVID-19 (**Fig. 1A**, **Fig. S2** and **Table S2**). Binding affinity is achieved through a combination of interface contact optimization and protein stability (**Fig. 3E**). While the RBD—ACE2 complex can be resolved at high-resolution by cryo-EM (*17, 23*), MD simulations provide orthogonal information about the interaction dynamics on a nanosecond timescale. In the case of CoVs, MD simulations reveal an exceptional versatility of viral receptor binding strategies (**Fig. 3E**). COVID-19 adopted a different strategy for achieving comparable affinity to SARS-2002: the interface of COVID-19 is significantly larger than that of SARS-2002 (1204Å vs. 998Å) with a remarkable number of interacting residues (ACE2: 30 vs. 24, **Fig. 1C**). In contrast, SARS-2002 is more flexible in its interaction with ACE2, interacting through fewer contacts that serve as “hot spots”. Therefore, we predict that SARS-2002 RBD neutralizing antibodies will not be effective for COVID-19. The failure of several of these antibodies to neutralize the binding of COVID-19 RBD to its receptor is consistent with our findings (*20, 23*). The fluctuation from high- to low-affinity conformations in SARS-2002 leads to an increased efficacy for inhibiting peptides (*24*) and high-affinity antibodies (*25*) compared to COVID-19. This implies a therapeutic challenge is attributed to the enhanced rigidity of the COVID-19 RBD relative to that of the SARS-2002.

The geometric and physicochemical properties of RBD—ACE2 interfaces resemble those of antibody-antigen interactions. In both cases the interface benefits from long loop plasticity, bulky aromatic side chains as anchoring sites, and the stabilization of the complex by distributed electrostatic interactions (*26*). Both COVID-19 and SARS-2002 interfaces contain long flexible loops and nine aromatic residues (Tyr, Trp, Phe) in the interface with ACE2 (**Fig. 2A**). Moreover, in the SARS designed variant (SARS-des (*11*)), the addition of an aromatic residue (L486F substitution) significantly improved the interaction scores and interface stability (**Fig. 3, B and D**). Our findings shed light on the accelerated evolution of spike protein binding to the ACE2 receptor similar to the rapid evolution along the antibody-antigen affinity maturation process.

## Acknowledgments

The authors gratefully acknowledge Barak Raveh for useful suggestions.

## Author contributions

EB, DS, and ML performed the calculations, analyzed the data, and prepared the manuscript

## Data and materials availability

The MD trajectories are available from ftp.huji.ac.il

## SUPPLEMENTARY MATERIALS

Materials and Methods Figs. S1–S3

Tables S1–S2

Movie S1 References (*27–29*)

## Materials and Methods

### Structural modeling

The structural model of the COVID-19 spike protein receptor binding domain (RBD) in complex with ACE2 was generated by comparative modeling using Modeller 9.18 (*27*) with the COVID-19 sequence (RefSeq: YP_009724390.1). We relied on the crystal structure of the spike protein receptor-binding domain from a SARS coronavirus designed human strain complexed with the human receptor ACE2 (PDB 3SCI, resolution 2.9Å) as a template for comparative modeling. The SARS-2002 spike protein RBD and HCoV-NL63 in complex with ACE2 were taken from PDB 2AJF (resolution 2.9Å) and 3KBH (resolution 3.3Å), respectively. Missing residues were added in MODELLER. MERS RBD structure was taken from the complex with the neutralizing antibody CDC2-C2 (PDB 6C6Z, resolution 2.1Å) and structurally aligned onto SARS-2002 RBD in complex with ACE2 receptor. The designed variant is from PDB 3SCI.

### Molecular dynamic simulations

The MD simulations were performed with GROMACS 2020 software (*28*) using the CHARMM36m force field (*29*). Each of the complexes was solvated in transferable intermolecular potential with 3 points (TIP3P) water molecules and ions were added to equalize the total system charge. The steepest descent algorithm was used for initial energy minimization until the system converged at Fmax < 1,000 kJ/(mol · nm). Then water and ions were allowed to equilibrate around the protein in a two-step equilibration process. The first part of equilibration was at a constant number of particles, volume, and temperature (NVT). The second part of equilibration was at a constant number of particles, pressure, and temperature (NPT). For both MD equilibration parts, positional restraints of k = 1,000 kJ/(mol nm^2^) were applied to heavy atoms of the protein, and the system was allowed to equilibrate at a reference temperature of 300 K, or reference pressure of 1 bar for 100 ps at a time step of 2 fs. Following equilibration, the production simulation duration was 100 nanoseconds with 2 fs time intervals. Altogether 10,000 frames were saved for the analysis at intervals of 10 ps. We superimposed several MD snapshots on the recently submitted to the PDB x-ray structure (6VW1, resolution 2.7Å) of COVID-19—ACE2 complex. The average RMSD over the interface Cα atoms is ~1Å. Interaction scores between the virus spike RBD and ACE2 were calculated for each frame of the trajectory using the SOAP statistical potential (*16*). In the interface contact analysis, a residue-residue contact was defined based on the inter-atomic distance, with a cutoff of 4Å.

---

**Fig. S1.**
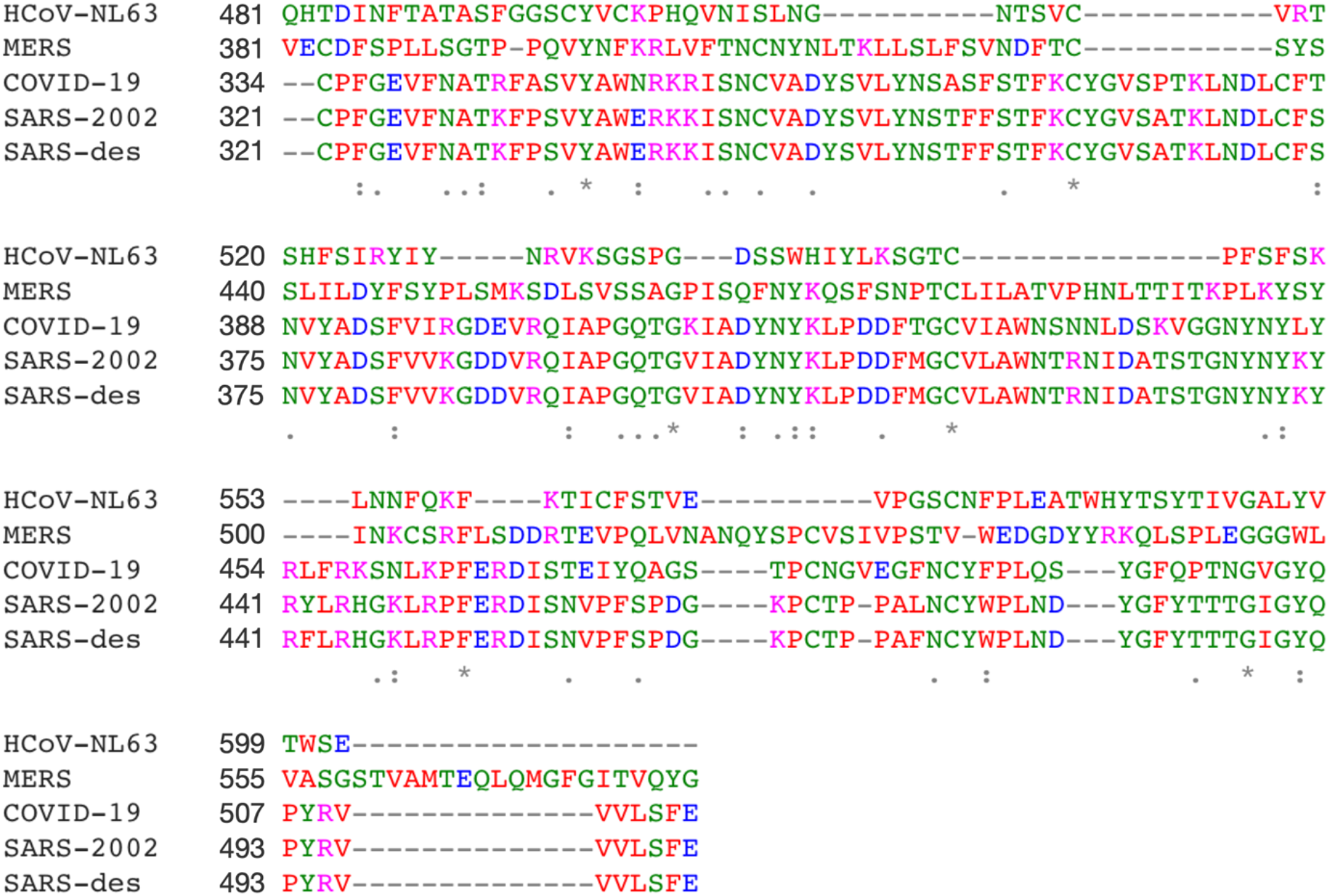
Multiple sequence alignment of the analyzed RBDs

**Fig. S2.**
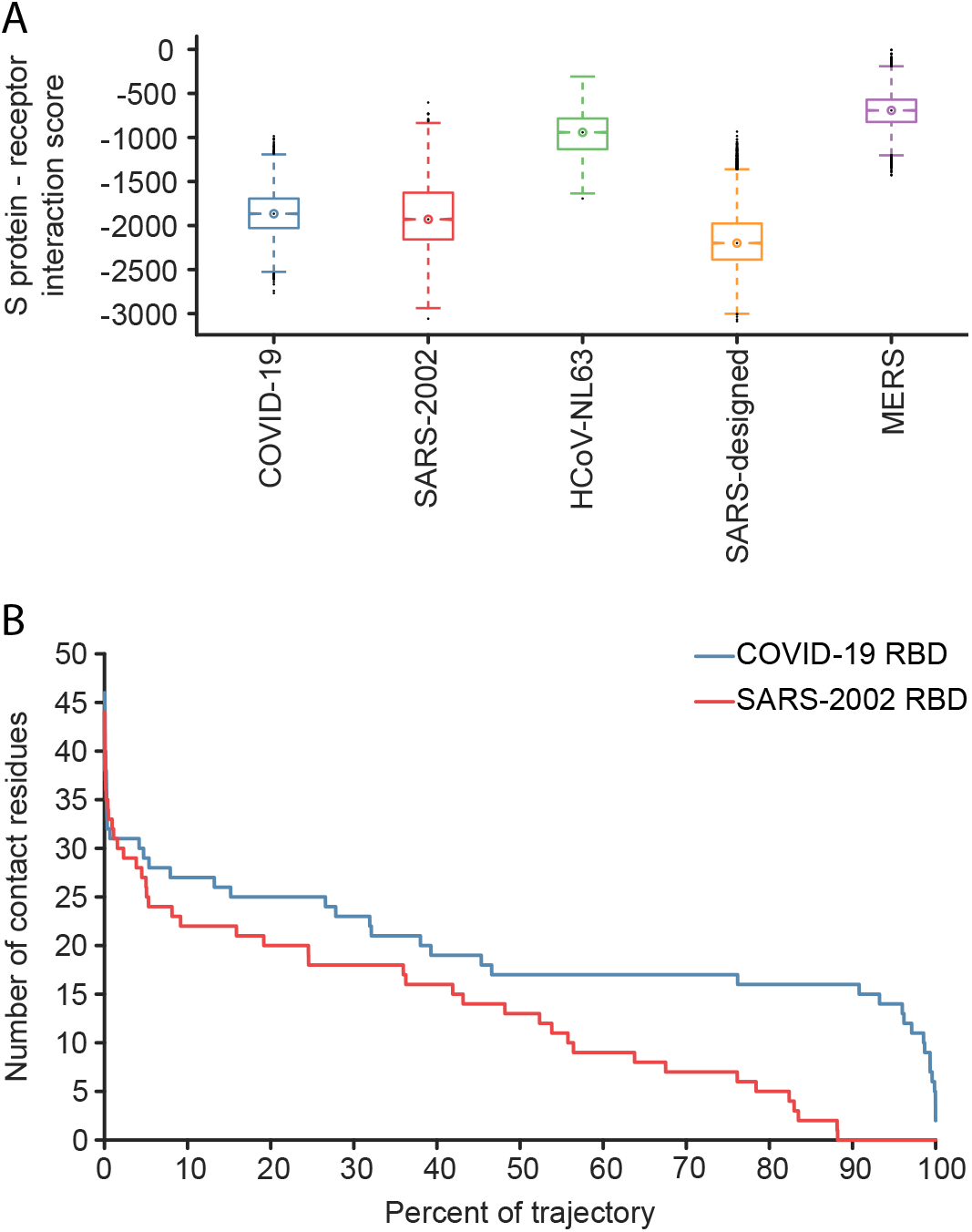
**(A)** Numbers of RBD residues in contact with ACE2 vs. the fraction of trajectory frames for COVID-19 (blue) and SARS-2002 (red). **(B)** Box plots of the SOAP interaction scores for the trajectories frames. The center point is the median score, while 50% of the scores are within the box. The whiskers extend to cover >99% of the scores.

**Fig. S3.**
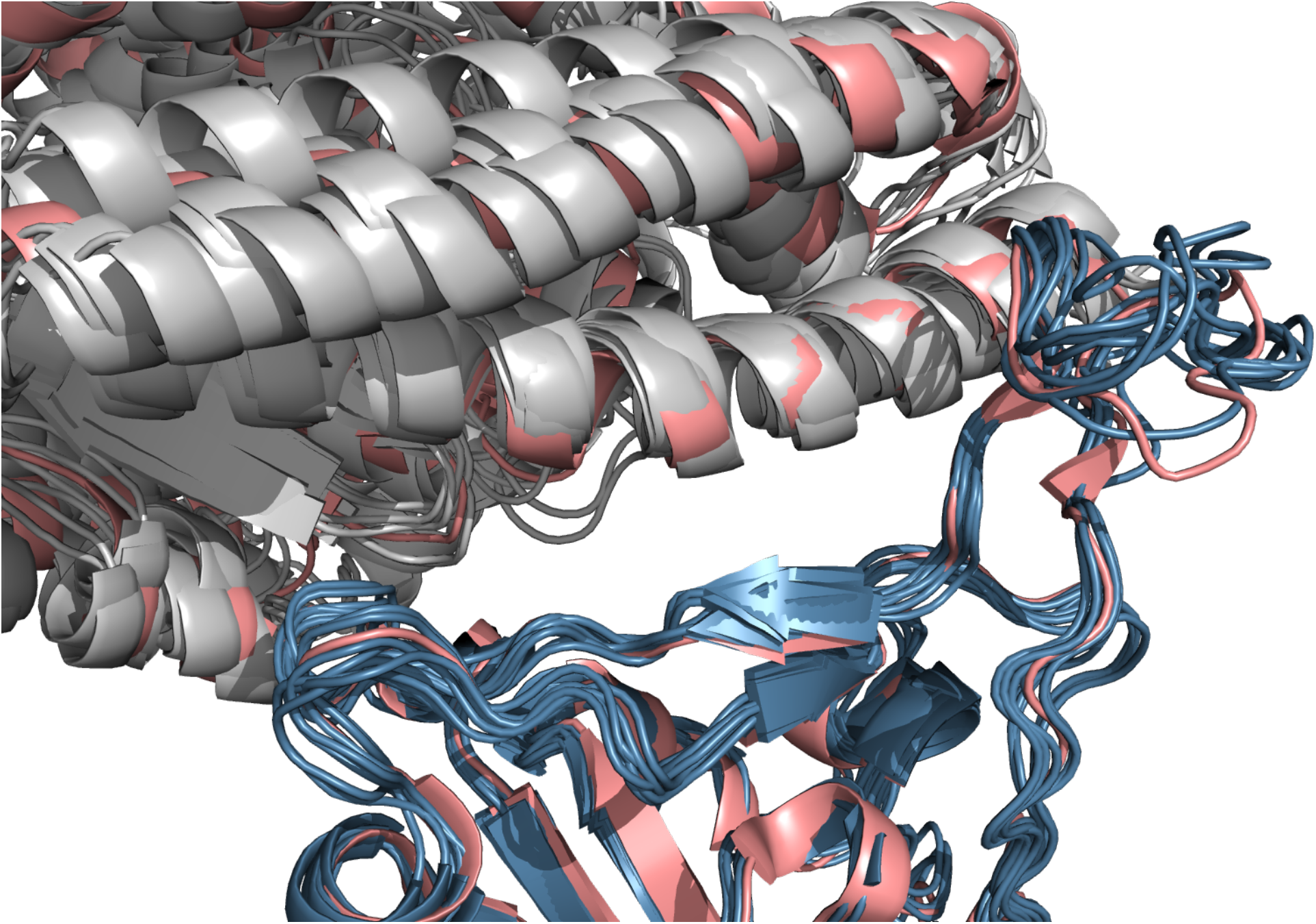
Comparison between the x-ray structure of COVID-19—ACE2 complex (PDB 6VW1, pink) and MD trajectory frames. The MD snapshots are colored gray and blue for ACE2 and COVID-19 RBD, respectively. The average RMSD over the interface Cα atoms is ~1Å.

**Table S1.** RBD interface residues that have substitutions between COVID-19 and SARS-2002

| COVID-19 | Residue number | SARS-2002 | Residue number |
| --- | --- | --- | --- |
| ARG | 403 | LYS | 390 |
| LYS | 417 | VAL | 404 |
| VAL | 445 | SER | 432 |
| GLY | 446 | THR | 433 |
| LEU | 455 | TYR | 442 |
| PHE | 456 | LEU | 443 |
| TYR | 473 | PHE | 460 |
| GLN | 474 | SER | 461 |
| ALA | 475 | PRO | 462 |
| GLY | 476 | ASP | 463 |
| SER | 477 | GLY | 464 |
| THR | 478 | LYS | 465 |
| ASN | 481 | THR | 468 |
| GLY | 482 | PRO | 469 |
| GLU | 484 | PRO | 470 |
| GLY | 485 | ALA | 471 |
| PHE | 486 | LEU | 472 |
| PHE | 490 | TRP | 476 |
| GLN | 493 | ASN | 479 |
| SER | 494 | ASP | 480 |
| GLN | 498 | TYR | 484 |
| PRO | 499 | THR | 485 |
| ASN | 501 | THR | 487 |
| VAL | 503 | ILE | 489 |

**Table S2.** RBD—ACE2 interface evaluated by several methods for analysis of protein-protein interactions

| | PRODIGY<br>$\Delta G$ (kcal mol<sup>-1</sup>) | PRODIGY<br>Kd 10<sup>-9</sup> (M) | FoldX<br>(kcal/mol<sup>-1</sup>) | SOAP<br>score |
| --- | --- | --- | --- | --- |
| COVID-19—ACE2 | -11.7 | 2.60 | -1.93 | -1865.9 |
| SARS-2002—ACE2 | -12.1 | 1.30 | -2.67 | -1929.5 |

**Movie S1.** Overlay of 50 random snapshots from the MD trajectories of COVID-19—ACE2, SARS-2002— ACE2, and HCoV-NL63—ACE2 complexes. For clarity only one copy of ACE2 is shown (gray), COVID-19, SARS-2002, and HCoV-NL63 are colored blue, red, and green, respectively.

## REFERENCES AND NOTES

1. P. A. Rota et al., Characterization of a novel coronavirus associated with severe acute respiratory syndrome. Science 300, 1394–1399 (2003).

2. L. E. Gralinski, V. D. Menachery, Return of the Coronavirus: 2019-nCoV. Viruses 12, 135 (2020).

3. S. R. Weiss, S. Navas-Martin, Coronavirus pathogenesis and the emerging pathogen severe acute respiratory syndrome coronavirus. Microbiol Mol Biol Rev 69, 635–664 (2005).

4. Z. Q. Zeng et al., Epidemiology and clinical characteristics of human coronaviruses OC43, 229E, NL63, and HKU1: a study of hospitalized children with acute respiratory tract infection in Guangzhou, China. Eur J Clin Microbiol Infect Dis 37, 363–369 (2018).

5. F. Wu et al., A new coronavirus associated with human respiratory disease in China. Nature, 1–5 (2020).

6. M. A. Tortorici et al., Structural basis for human coronavirus attachment to sialic acid receptors. Nature structural & molecular biology 26, 481–489 (2019).

7. K. Kuba et al., A crucial role of angiotensin converting enzyme 2 (ACE2) in SARS coronavirus– induced lung injury. Nature medicine 11, 875–879 (2005).

8. H. Hofmann et al., Susceptibility to SARS coronavirus S protein-driven infection correlates with expression of angiotensin converting enzyme 2 and infection can be blocked by soluble receptor. Biochemical and biophysical research communications 319, 1216–1221 (2004).

9. E. C. Mossel et al., Exogenous ACE2 expression allows refractory cell lines to support severe acute respiratory syndrome coronavirus replication. J Virol 79, 3846–3850 (2005).

10. I. Hamming et al., Tissue distribution of ACE2 protein, the functional receptor for SARS coronavirus. A first step in understanding SARS pathogenesis. The Journal of Pathology: A Journal of the Pathological Society of Great Britain and Ireland 203, 631–637 (2004).

11. Y. Wan, J. Shang, R. Graham, R. S. Baric, F. Li, Receptor recognition by novel coronavirus from Wuhan: An analysis based on decade-long structural studies of SARS. J Virol (2020).

12. H. Hofmann et al., Human coronavirus NL63 employs the severe acute respiratory syndrome coronavirus receptor for cellular entry. Proc Natl Acad Sci U S A 102, 7988–7993 (2005).

13. W. Li et al., The S proteins of human coronavirus NL63 and severe acute respiratory syndrome coronavirus bind overlapping regions of ACE2. Virology 367, 367–374 (2007).

14. N. Wang et al., Structure of MERS-CoV spike receptor-binding domain complexed with human receptor DPP4. Cell research 23, 986 (2013).

15. Y. Zhang, J. Skolnick, TM-align: a protein structure alignment algorithm based on the TM-score. Nucleic acids research 33, 2302–2309 (2005).

16. G. Q. Dong, H. Fan, D. Schneidman-Duhovny, B. Webb, A. Sali, Optimized atomic statistical potentials: assessment of protein interfaces and loops. Bioinformatics 29, 3158–3166 (2013).

17. R. Yan, Y. Zhang, Y. Guo, L. Xia, Q. Zhou, Structural basis for the recognition of the 2019-nCoV by human ACE2. bioRxiv (2020).

18. Y. Zhang, N. Zheng, Y. Zhong, Computational characterization and design of SARS coronavirus receptor recognition and antibody neutralization. Computational biology and chemistry 31, 129–133 (2007).

19. W. Li et al., Receptor and viral determinants of SARS-coronavirus adaptation to human ACE2. EMBO J 24, 1634–1643 (2005).

20. X. Tian et al., Potent binding of 2019 novel coronavirus spike protein by a SARS coronavirus-specific human monoclonal antibody. Emerg Microbes Infect 9, 382–385 (2020).

21. K. Wu, W. Li, G. Peng, F. Li, Crystal structure of NL63 respiratory coronavirus receptor-binding domain complexed with its human receptor. Proceedings of the National Academy of Sciences 106, 19970–19974 (2009).

22. A. C. Walls et al., Structure, function and antigenicity of the SARS-CoV-2 spike glycoprotein. bioRxiv (2020).

23. D. Wrapp et al., Cryo-EM structure of the 2019-nCoV spike in the prefusion conformation. Science (2020).

24. A.-W. Struck, M. Axmann, S. Pfefferle, C. Drosten, B. Meyer, A hexapeptide of the receptor-binding domain of SARS corona virus spike protein blocks viral entry into host cells via the human receptor ACE2. Antiviral research 94, 288–296 (2012).

25. L. Du et al., The spike protein of SARS-CoV--a target for vaccine and therapeutic development. Nat Rev Microbiol 7, 226–236 (2009).

26. V. Kunik, Y. Ofran, The indistinguishability of epitopes from protein surface is explained by the distinct binding preferences of each of the six antigen-binding loops. Protein Eng Des Sel 26, 599–609 (2013).

27. B. Webb, A. Sali, Comparative protein structure modeling using MODELLER. Current protocols in bioinformatics 47, 5.6. 1–5.6. 32 (2014).

28. S. Pronk et al., GROMACS 4.5: a high-throughput and highly parallel open source molecular simulation toolkit. Bioinformatics 29, 845–854 (2013).

29. J. Huang et al., CHARMM36m: an improved force field for folded and intrinsically disordered proteins. Nature methods 14, 71–73 (2017).

